# Functional and compositional changes in the fecal microbiome of a shorebird during pre-migratory weight gain

**DOI:** 10.1101/2022.06.30.498374

**Authors:** Kirsten Grond, Artemis S. Louyakis, Sarah M. Hird

**Author notes:** These authors contributed equally to this work (Kirsten Grond, Sarah M. Hird).

## Abstract

Shorebirds migrate long distances twice annually, which requires intense physiological and morphological adaptations, including the ability to rapidly gain weight via fat deposition at stopover locations. The role of the microbiome in weight gain in avian hosts is unresolved, but there is substantial evidence to support the hypothesis that the microbiome is involved with host weight from mammalian microbiome literature. Here, we collected 100 fecal samples of Ruddy Turnstones to investigate microbiome composition and function during stopover weight gain in Delaware Bay, USA. Using 16S rRNA sequencing on 90 of these samples and metatranscriptomic sequencing on 22, we show that taxonomic composition of the microbiome shifts during weight gain, as do functional aspects of the metatranscriptome. We identified ten genes that are associated with weight class and polyunsaturated fatty acid biosynthesis in the microbiota is significantly increasing as birds gain weight. Our results support that the microbiome is a dynamic feature of host biology that interacts with both the host and the environment and may be involved in the rapid weight gain of shorebirds.

## INTRODUCTION

Migratory birds go through a myriad of physiological changes throughout their annual cycle. The most extreme changes are associated with the act of migration [1, 2]. Shorebirds (Order: Charadriiformes) undertake migrations of thousands of kilometers, twice a year, between their breeding grounds and non-breeding grounds (Fig. 1). To prepare for migration, shorebirds absorb part of their digestive tract, which is unused during flight, to reduce body weight. They also increase the size of their pectoral muscles to maximize flight performance [3, 4]. Prior to migration, shorebirds rapidly gain weight to fuel their flights, often almost doubling their body mass in as little as 14 days [5]. This rapid weight gain is due to a short period of extreme foraging behavior, called hyperphagia. Birds increase their food intake by 20-40% and can accomplish a 7% mass gain per day [6, 7]. The weight is largely comprised of fat, which provides the most efficient fuel to complete their migrations. The causes and consequences of these physiological changes, including that they relate to a complex and demanding vertebrate behavior, continue to be explained. One major question with both basic and applied implications is the role of the microbiome.

**Figure 1.**
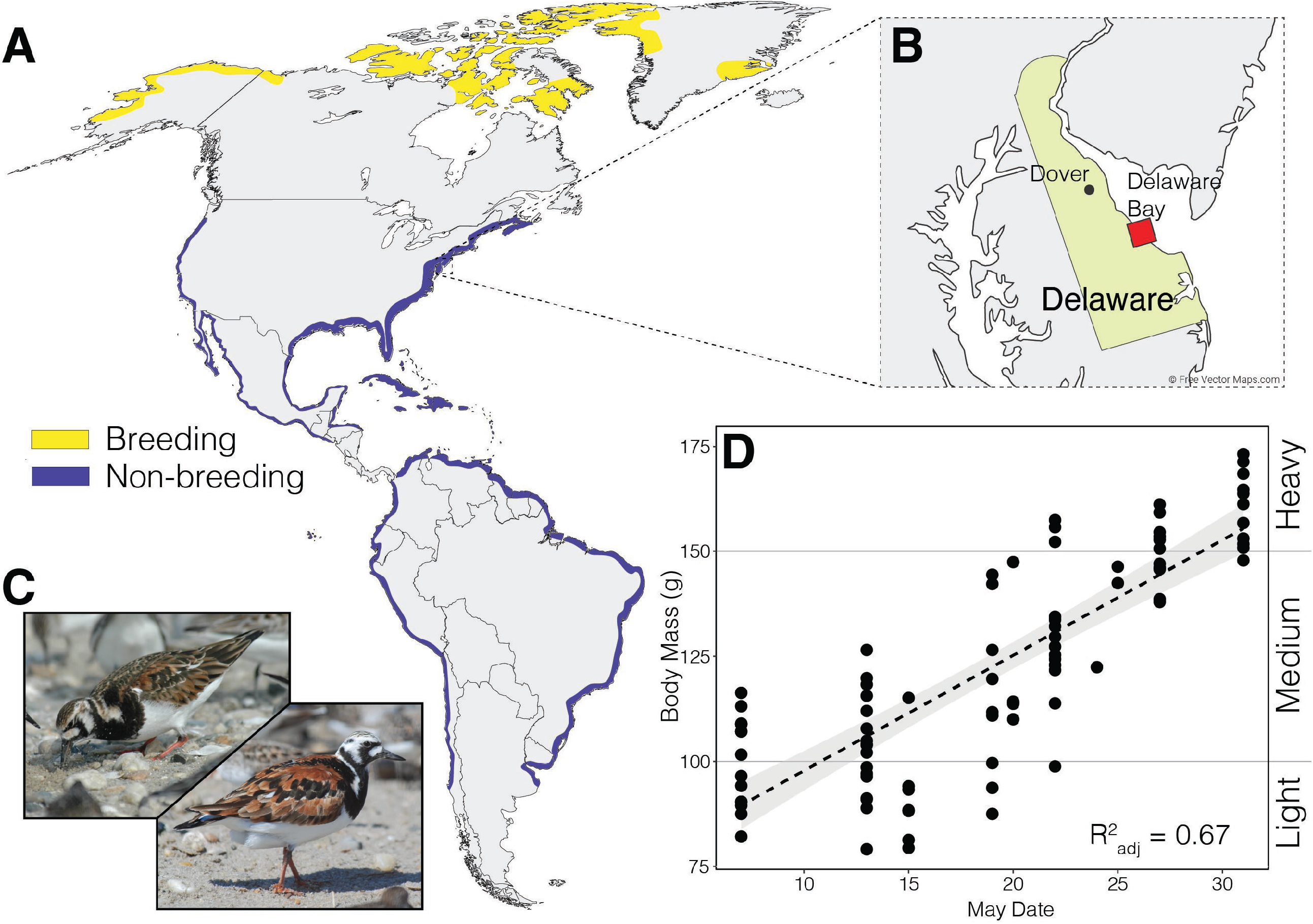
Ruddy Turnstone attributes. (A) Distribution of North, Central and South American breeding and non-breeding grounds. (B) Sampling site (Delaware Bay, DE). (C) Female (left) and male (right) breeding plumage, photos (C) Gregory Breese, US Fish and Wildlife Service. (D) Weight distribution of 91 Ruddy Turnstones sampled during stopover in 2018, linear regression line with region of standard error highlighted in gray (adjusted R^2^ = 0.67).

A vertebrate’s microbiome – the microorganisms that live on and inside a host – is intimately involved in many aspects of vertebrate biology, including development, immunity, behavior and digestion (reviewed in [8]). The microbiome is associated with weight gain, with an applied focus on humans and model organisms [9, 10]. Bacteria within the phylum *Firmicutes* has been associated with obesity when paired with a high fat diet ([11] but see [12]); conversely, other bacteria (notably *Bacteroidetes*) are associated with lean or normal weight [13–15]. Obese mice with microbiomes rich in *Firmicutes* extract more energy from given amount of food than lean mice with microbiomes relatively lower in *Firmicutes* [13]. Strong mechanistic links between the microbiome and fat deposition involve bacterial metabolites, host gene regulation, and lipogenesis [16]. The gut-microbiome-brain axis posits additional ways that the microbiome can influence weight in hosts, through food-seeking behavior, appetite, taste and food preferences (reviewed in [17]). Similar to the natural, high-fat state of pre-migratory shorebirds, many species of mammals rapidly gain weight before hibernation, which sustains them through periods of low food intake. For example, arctic ground squirrels (*Urocitellus parryii)* hibernate for 6-9 months and rely completely on the fat mass accumulated during their active season. Although the role of the microbiome in fattening is unknown in this species, the microbiome was recently shown to be involved in maintaining lean mass during hibernation [18]. In another hibernator, the brown bear (*Ursus arctos*), the microbiome undergoes annual changes. During the summer, while the bears are gaining weight, the microbiome is relatively higher in Firmicutes, which can increase fat deposition under controlled conditions when transplanted into mice [19].

There is a clear link between the microbiome and weight in birds. Antibiotics were widely administered for decades for the purposes of increasing growth rate and weight gain in livestock and poultry [20]. As the side effects of these widespread practices became apparent, the industry moved away from antibiotics and has turned to probiotics to manipulate livestock and poultry microbiomes, which can also positively impact weight in birds (reviewed in [21]). In wild shorebirds, the increased food input combined with internal physiological shifts raises the question of how the microbiome affects and is affected by these changes. The taxa within the microbiome are frequently decoupled from the functional potential of the community (e.g., [15]) with dramatic differences in variance of these two metrics and questions remain about the relative roles of each in community assembly and host-microbe interactions. Understanding how taxonomic composition and microbial function are related to physiological changes may indicate the mechanisms used in adjusting to changing environmental conditions.

The compositional dynamics of the microbial taxa in the gut microbiome of shorebirds has been studied using 16S rRNA gene amplicon sequencing at different stages of the annual cycle. Geographic location is significantly correlated to the taxonomic composition of the microbiome across multiple host species [22] but in highly similar environments, bacterial community structure can be host species specific [23]. After hatch, shorebird gut communities grow exponentially for two days and then stabilize [24]; their long-distance migrations impact the composition of the microbiome [25] but are also resistant to many potentially horizontally transferred bacteria [26]. These 16S rRNA-based studies have robustly characterized taxonomic dynamics. However, bacterial communities can assemble based on function [27] and phylogenetic and functional diversity can reveal unique aspects of microbial communities [28] Metatranscriptomics complements 16S rRNA amplicon data by extracting the total RNA in a sample and enriching for and sequencing the mRNA to identify the recently transcribed genes. Transcriptomics can reveal patterns at a relatively fine scale, resulting in a deep characterization of the active processes, such as pathway utilization and unexpected gene activity. A comprehensive investigation into the taxonomic composition and functional expression of the microbiome in wild migrating birds will provide insight into how microorganisms relate to a physiologically critical period in a vertebrate.

We investigated the Ruddy Turnstone (*Arenaria interpres*) fecal microbiome, collected in Delaware Bay at different stages of fattening during the spring-migration staging period, using both 16S rRNA gene amplicon and metatranscriptome sequencing. Ruddy Turnstones stop in Delaware Bay for approximately 2 weeks during the month of May on their northwards migration and double their body mass during this time period by intensive foraging on the eggs of the Horseshoe Crab, *Limulus polyphemus* [29]. The primary aims of our study were: 1) compare taxonomic and functional patterns in shorebird microbiomes during pre-migratory fattening, 2) identify significantly differentially expressed microbial genes to elucidate pathways important to changing host weight and potential variation between sexes.

## METHODS

### Sample Collection

Fecal samples were collected from 100 Ruddy Turnstones (*Arenaria interpres*) from 7-31 May 2018 at three beaches in Delaware Bay, DE (Fig. 1). Birds were captured using cannon nets as part of the Delaware Shorebird Project, a program from the Delaware Department of Natural Resources and Environmental Control. Upon capture, birds were placed in individual boxes lined with 10% bleach-sterilized trays for up to 10 minutes. A mesh platform above the tray avoided contamination of fecal samples by the birds’ feet; see [30] for detailed sampling description. Fecal samples were preserved in DNA/RNA shield (Zymo Research, Irvine CA) upon collection, and frozen at -20°C within two hours of capture. After sample collection, weight and biometric measurements (wing length, head and bill length) were collected and birds were sexed and aged based on plumage characteristics (Fig. 1C).

Samples were sorted into three weight classes to increase sample size for statistical analysis. Birds were classified as light (<100g), medium (100-150g), and heavy (>150g). The medium weight category starts at 100g, as this is the average weight of Turnstones during the wintering period [31]. Many (if not all) birds, when they first land at Delaware Bay, are below the average wintering period weight and thus classified as “light” weight. “Heavy” was defined as a 50% increase above the wintering period average. Because all birds lose weight during migratory flight and gain weight during stopover, the weight categories are approximate indicators of how long they have been at the stopover location and how soon they may begin the next leg of their migration.

### DNA Extraction & Sequencing

DNA and RNA were extracted simultaneously using the ZymoBIOMIC™ DNA/RNA Miniprep kit (Zymo Research, Irvine CA), following the parallel extraction protocol. Extracted RNA and DNA were stored at -80°C until sequencing.

For the 16S rRNA gene sequencing, the V4 region of the 16S rRNA gene was PCR amplified and sequenced at the University of Connecticut Microbial Analysis, Resources, and Services facility, following the standard operating procedure. The V4 region of the 16S rRNA gene was sequenced at the UConn Microbial Analysis, Resources, and Services facility. Quant-iT PicoGreen kit was used to quantify DNA concentrations, and 30 ng of extracted DNA was used as template to amplify the V4 region of the 16S rRNA gene. V4 primers (515F and 806R) with Illumina adapters and dual barcodes were used for amplification [32, 33]. PCR conditions consisted of 95°C for 3.5 min, 30 cycles of 30 s at 95.0°C, 30 s at 50.0°C and 90 s at 72.0°C, followed by final extension at 72.0°C for 10 min. PCR products were normalized based on the concentration of DNA from 250-400 bp and pooled. Pooled PCR products were cleaned using the Mag-Bind RxnPure Plus (Omega Bio-tek) according to the manufacturer’s protocol, and the cleaned pool was sequenced on the MiSeq using v2 2×250 base pair kit (Illumina, Inc, San Diego, CA).

For the metatranscriptomes, total RNA was quantified, and purity ratios determined for each sample using the NanoDrop 2000 spectrophotometer (Thermo Fisher Scientific, Waltham, MA, USA). To assess RNA quality, total RNA was analyzed on the Agilent TapeStation 4200 (Agilent Technologies, Santa Clara, CA, USA) using the RNA High Sensitivity assay following the manufacturers protocol. Ribosomal Integrity Numbers (RIN) were recorded for each sample. Total RNA samples (300ng of Qubit quantified total RNA input) were prepared for prokaryotic transcriptome sequencing by first ribodepleting bacterial ribosomal RNA using the RiboMinus Transcriptome Isolation Kit, Bacteria (ThermoFisher Scientific, Waltham, MA, USA).

Ribodepletion efficiency was analyzed prior to the start of library preparation on the Agilent TapeStation 4200 (Agilent Technologies, Santa Clara, CA, USA) using the RNA High Sensitivity assay following the manufacturers protocol. Efficient ribodepletion is supported by the disappearance of the 16S and 23S ribosomal RNA peaks (∼1,000 nt and ∼2,000 nt, respectively), with the sample’s electropherogram trace now showing a smear of shorter molecules (< 1,000nt).

Purified ribodepleted RNA underwent library preparation using the Illumina TruSeq Stranded mRNA Sample Preparation kit following the manufacturer’s protocol modification for purified mRNA as input (Illumina, San Diego, CA, USA). Libraries were validated for length and adapter dimer removal using the Agilent TapeStation 4200 D1000 High Sensitivity assay (Agilent Technologies, Santa Clara, CA, USA) then quantified and normalized using the dsDNA High Sensitivity Assay for Qubit 3.0 (Life Technologies, Carlsbad, CA, USA). Sample libraries of sufficient quality were sequenced (Illumina MiSeq; paired end 2 × 75bp read length) with a sequencing depth targeted at 7-10M total paired end reads/sample at the Center for Genome Innovation at the University of Connecticut.

### Sequence quality control, assembly, annotation, and mapping

For the 16S rRNA gene amplicon data, during standard Illumina demultiplexing, sequences were quality checked and trimmed to remove adaptors and barcodes. The DADA2 (v. 3.11) pipeline in R (v3.6.0) was used to quality control and process the reads [34, 35], low quality read areas were removed following the DADA2 default parameters. Following assessment of error rates, paired-end sequences were merged, and potentially chimeric sequences removed. All unique sequences at greater than 1x abundance were then labeled as amplicon sequence variants, or ASVs, for taxonomic analysis. Sequences were assigned to taxonomy using RDP’s Naïve Bayesian Classifier with the Silva (v. 132) reference database [36, 37]. Sequences identified as chloroplast or mitochondrial sequences were removed from the dataset. A multiple-sequence alignment was performed using the *DECIPHER* (v. 2.0) package [38], and a phylogenetic tree was constructed with the package *phangorn* (v2.4.0; [39]). Likely sequence contaminants were identified and removed using the *decontam* (v1.4.0; [40]) package using the negative control samples as contaminants.

The metatranscriptome sequences were trimmed using Trimmomatic (v0.35; [41]) with a threshold of Q5 and rRNA was removed using SortMeRNA [42]). The remaining sequences were de novo assembled using Trinity (v2.2.0; [43]) with the following parameters: fastq assembly (left read file contained forward and unpaired reads), minimum contig length of 75 bp and normalized reads. Alignment was completed using Bowtie2 (v2.2.9; [44]) and RSEM (v1.2.7, [45]) estimation was used for counts of sample replicates. RSEM estimates were rounded to nearest integer, length corrected (transcripts per million method), trimmed mean of M-values (TMM) adjusted for normalized expression values (EdgeR v3.16.5; [46]), and batch corrected with ARSyNseq (in NOISeq,[47, 48]). The metatranscriptome assembly was annotated using Trinotate (v3.0.1) using the complete pipeline (http://trinotate.github.io).

### Statistical methods

A linear regression was used to confirm weight gain of Ruddy Turnstone over the stopover period. To assess microbiome taxonomic compositional change with weight gain, alpha and beta diversity measures were calculated for the 16S rRNA gene amplicons. For alpha diversity analyses, samples were rarefied to 10 995 sequences, which was the lowest sequence coverage of our samples. Two alpha diversity measures were calculated using *phyloseq* (v1.28.0; [49]): the observed number of ASVs and Shannon’s Diversity Index [50]. Statistical significances for differences in alpha diversity for sites, weight classes, and sexes were calculated using analysis of variance (ANOVA) testing. For beta diversity, three distance metrics were calculated to describe differences between samples: Bray-Curtis dissimilarity, weighted UniFrac and unweighted UniFrac [51]. These distance matrices were used for non-metric multidimensional scaling (NMDS). The relative contributions to the variation in microbiome composition of three variables (weight class, sex, and sampling site) were calculated using permutational multivariate analysis of variance (PERMANOVA) with the adonis2 function from the *vegan* package (v2.5.6; [52]). We tested for homogeneity of variance among weight classes and sexes using the betadisper function, also in *vegan*. Differential abundance in taxa between weight classes was conducted using *DESeq2* package in R (v1.24.0; [53]). Significance was set at α=0.001.

We conducted a Principal Components Analysis on the TMM normalized expression counts and identified significance and relative contributions of weight class and sex in functional composition. A pairwise PERMANOVA using the *pairwise*.*perm*.*manova* function in the *RVAideMemoire* package (v. 0.9.75; [54]) with 1 000 permutations was used to assess significance between weight classes.

Differential gene expression between weight classes and sexes was conducted using *NOISeq* package in R (v1.24.0;.[47]) with significance set to at α=0.001. Volcano plots were constructed from NOISeq data using the *EnhancedVolcano* (v1.2.0; [55]) and *ggplot2* (v3.3.0; [56]) packages. To identify similar patterns of expression among differentially expressed genes, a cluster analysis using the *DESeq2* (v1.24.0; [53]) and the *DEGreport* [57] packages was conducted. Likelihood Ratio Testing (α_adj_=0.001/0.01/0.05) identified differences in expression across all weight classes and identified gene clusters across groups using the degPatterns function from *DEGreport*.

Because Ruddy Turnstones’ main activity during stopover is acquiring fat mass, we focused on genes and pathways related to lipid metabolism. A variance stabilizing transformation was applied to count matrices using the vst function in *DeSeq2*. To assess the relationship between body weight of Ruddy Turnstones and specific expressed pathways, normalized expression plots were constructed for biosynthesis of all Fatty Acids (FAs), and three essential Polyunsaturated Fatty Acids (PUFAs): Arachidonic acid, Linoleic acid, and alpha-Linolenic acid.

## RESULTS

Ruddy Turnstones consistently gained weight over the 2018 mid-migration stopover at Delaware Bay (Fig. 1; Linear Regression Model, F_1,89_=182.8, adj. R^2^=0.67, *p<*0.001). Body mass significantly differed among our three sampling sites (ANOVA: F_2,88_=78.51, *p<*0.001), with the lowest bird weights at Swains and the highest weights at Back North (Fig. S1).

From the 100 fecal samples collected from Ruddy Turnstones, 90 samples were of sufficient post-extraction quality for 16S rRNA gene sequencing. We sampled 45 female (F) and 41 male (M) Turnstones. Four individuals could not reliably be assigned a sex and are referred to as unknown (U). After quality control, we retained 3 700 042 high quality sequences across the 90 samples with an average of 40 660 ± 1 733 SE sequences per sample.

### Taxonomic Composition and Diversity Using 16S rRNA

#### Alpha diversity

Shannon’s diversity index (“Shannon”) significantly differed among weight classes (Fig. 2A, ANOVA: Shannon F_2,88_=5.648, *p=*0.005). Light and medium weight birds differed significantly from heavy birds, but not from each other (TukeyHSD: <100g - >150g, *p=*0.005; 100-150g - >150g, *p=*0.032; <100g – 100-150g, *p=*0.876). Observed number of ASVs did not significantly differ between weight classes (ANOVA: Observed F_2,88_=2.967, *p=*0.057). Alpha diversity significantly differed among sampling sites (ANOVA: Shannon F_2,88_=6.449, *p=*0.002; Observed F_2,88_=3.335, *p=*0.040), but did not differ between sexes (ANOVA: Shannon F_1,88_=0.136, *p=*0.873; Observed F_1,88_ =0.493, *p=*0.612).

**Figure 2.**
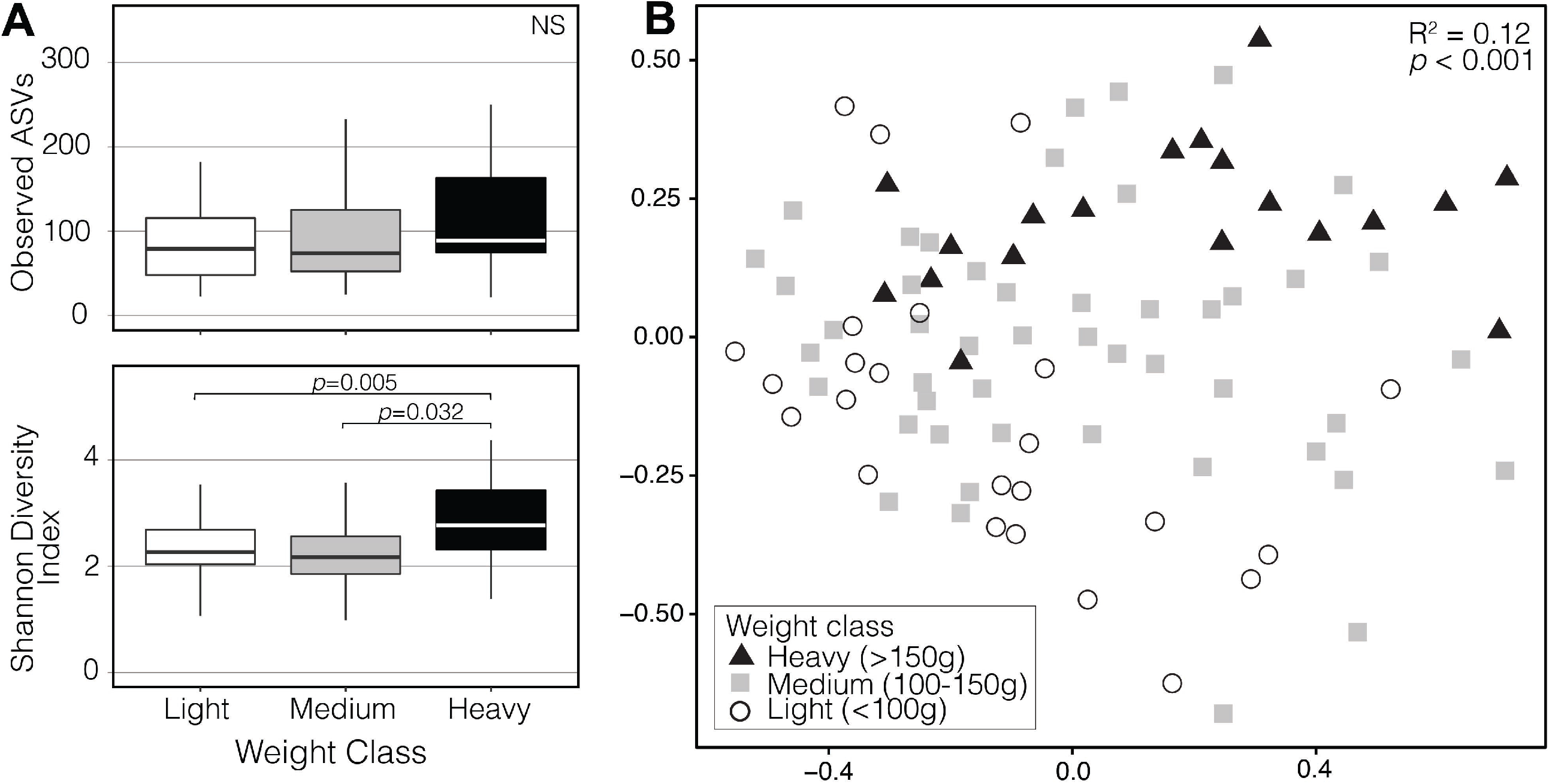
Alpha diversity of the three weight classes (*p*-values < 0.005 shown). (B) Non-metric Multidimensional Scaling ordination constructed from a Bray-Curtis matrix of 16S rRNA gene communities. Shapes and colors represent the three weight classes (black/triangle = Heavy, gray/square = Medium, white/circle = Light).

#### Beta diversity

The NMDS plot showed clustering in microbiome communities by weight class and showed directional change from light to heavy birds; medium weight class birds appeared to be located in between the light and heavy weight classes (Fig. 2B). Weight class (R^2^=12%) and sampling site (R^2^=3.6%) were significantly associated with variation in microbiome composition (*p<*0.001). Sex of the birds was not significantly correlated with microbiome composition (PERMANOVA: F_2,84_=0.62, R^2^=0.014, *p=*0.994). Homogeneity of variance (beta dispersion) did not significantly differ among weight classes (Permutest: F_8,84_= 0.75, *p=*0.668), sexes (Permutest: F_2,84_= 0.50, *p=*0.612), or sampling sites (Permutest: F_2,88_= 1.18, *p=*0.330).

#### Community Composition

Twenty-six phyla were identified across all samples; five phyla comprised 97.3% of all sequences. The dominant phylum was Fusobacteria (40.7%), followed by Proteobacteria (26.7%), Firmicutes (18.9%), Bacteroidetes (5.8%), and Tenericutes (5.2%; Fig. S1). The 784 genera detected contained 4 463 ASVs. The Fusobacteria phylum was dominated by two genera, *Fusobacterium* and *Cetobacterium*, which together comprised >99.9% of sequences within this phylum. *Helicobacter* and *Campylobacter* were the dominant genera within the Proteobacteria, *Catellicoccus* within the Firmicutes, and *Bacteroides* and *Flavobacterium* within the Bacteroidetes. Within the most abundant genera, only *Vibrio* and *Flavobacterium* were significantly different between any weight classes (Fig. 3; full statistical tests in Supplemental Table S1).

**Figure 3.**
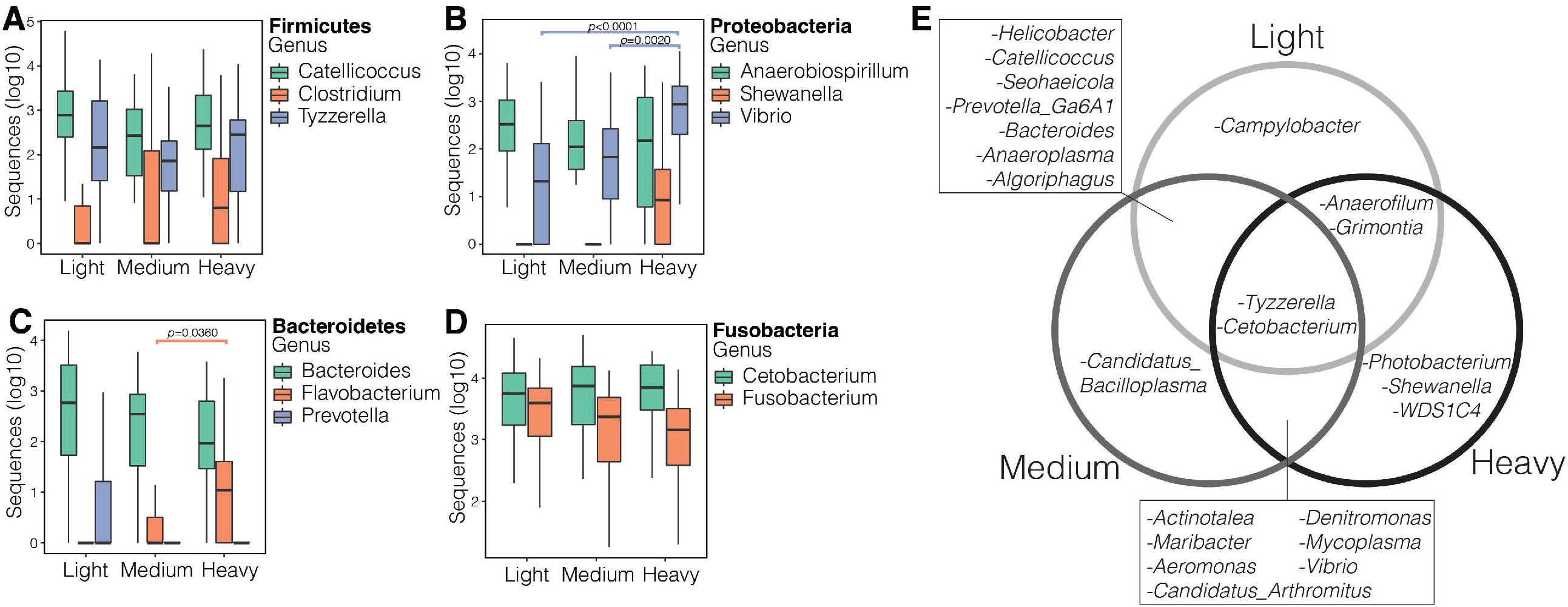
Relative abundance of the dominant genera within the four most abundant phyla (A) Firmicutes, (B) Proteobacteria, (C) Bacteroidetes, (D) Fusobacteria. Genera are separated by weight on the x-axis; all significant (*p*<0.05) changes within the genera across weight classes are noted with respective p-values. (E) Genera detected by DESeq2 analysis to be differentially abundant in two pairwise comparisons (light v medium, medium v heavy, light v heavy). NOTE: Because some genera had more than one ASV differentially expressed, and those ASVs might have been present in both categories, it’s possible to have genera overexpressed in all three categories.

#### Differential Abundance of Taxa

Pairwise tests identified many genera associated with weight class (Supplemental Table S2). Those that were consistent across both pairwise tests (e.g., taxa significantly associated with light weight birds in both the light against medium and light against heavy comparisons) revealed 13 differentially abundant genera. Since some of these genera contained more than one differentially expressed ASV, two genera were differentially expressed in all three weight categories: *Tyzzerella* and *Cetobacterium* (Fig. 3E). In the light birds, only *Campylobacter* was differentially expressed in both its pairwise comparisons, in the medium birds, only *“Candidatus Bacilloplasma*” and in the heavy birds, *Photobacterium, Shewanella* and *WDS1C4*. Furthermore, *Vibrio* was overexpressed in the medium and heavy birds, when compared to the light. *Helicobacter, Catellicoccus, Seohaeicola* and *Prevotella Ga6A1 group* were significantly higher in the light and medium birds when compared to the heavy and *Grimontia* was significantly higher in the light and heavy birds when compared to medium.

### Functional Dynamics of the Metatranscriptome

After ribodepletion and library preparation, 22 out of 40 initially selected RNA samples were suitable for metatranscriptome sequencing, including 14 males and eight females in the following weight class sample sizes: light (N=7), medium (N=9), and heavy (N=5).

We detected a significant difference in the functional gene community (based on KEGG IDs) among the three weight classes (Fig. 4, PerMANOVA: F_2,19_=2.78, R^2^=0.227, *p=*0.02), but not between sexes (PerMANOVA: F_1,20_=1.56, R^2^=0.027, *p=*0.605). Light birds and medium birds differed significantly from each other (*p*_adj_=0.018), but no difference in functional community was detected between light birds and heavy birds (*p*_adj_=0.366) or between medium birds and heavy birds (*p*_adj_=0.226). PCA beta dispersion differed among weight classes (ANOVA: F_2,19_=5.10, *p=*0.017), which was driven by a significant difference between light birds and heavy birds (TukeyHSD: *p*_adj_=0.020). Beta dispersion did not differ between light birds and medium birds (TukeyHSD: *p*_adj_=0.074) nor between medium birds and heavy birds (TukeyHSD: *p*_adj_=0.588).

**Figure 4.**
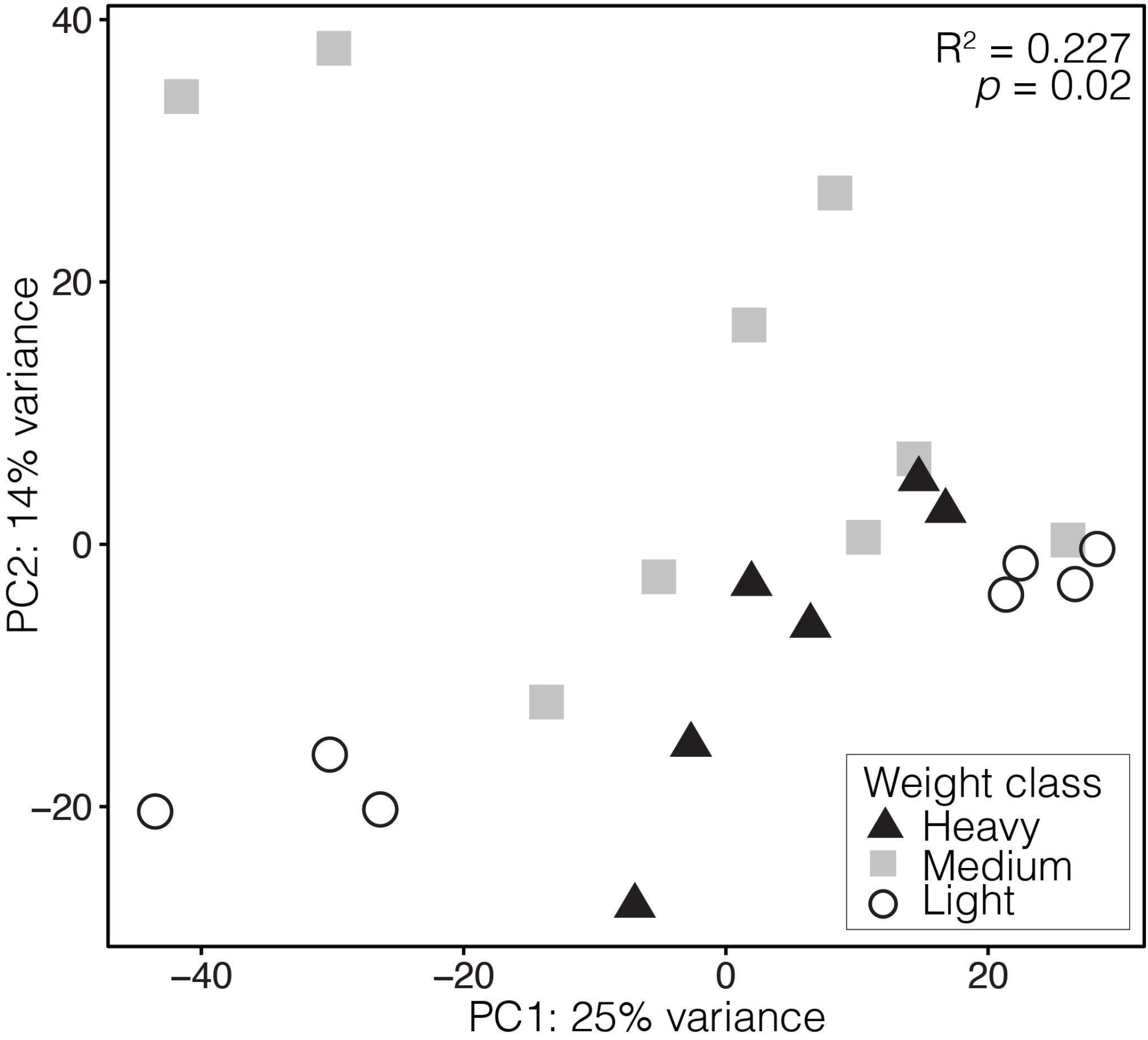
Principal components analysis of fecal metatranscriptomes collected from Ruddy Turnstones of different weights in Delaware Bay. Transcript counts were log-transformed, and colors/shapes represent the three bird weight classes.

#### Differential expression

Several KEGG objects were differentially expressed among weight classes (light vs. medium, N=7; light vs. heavy, N=4; medium vs. heavy, N=1; Fig. 5, Supplemental Table S3). One, K06422, was differentially expressed between female and male individuals, as well as in light birds. K06422 is an unclassified gene that is active in small acid-soluble spore protein E (*sspE*) production during cell growth.

**Figure 5.**
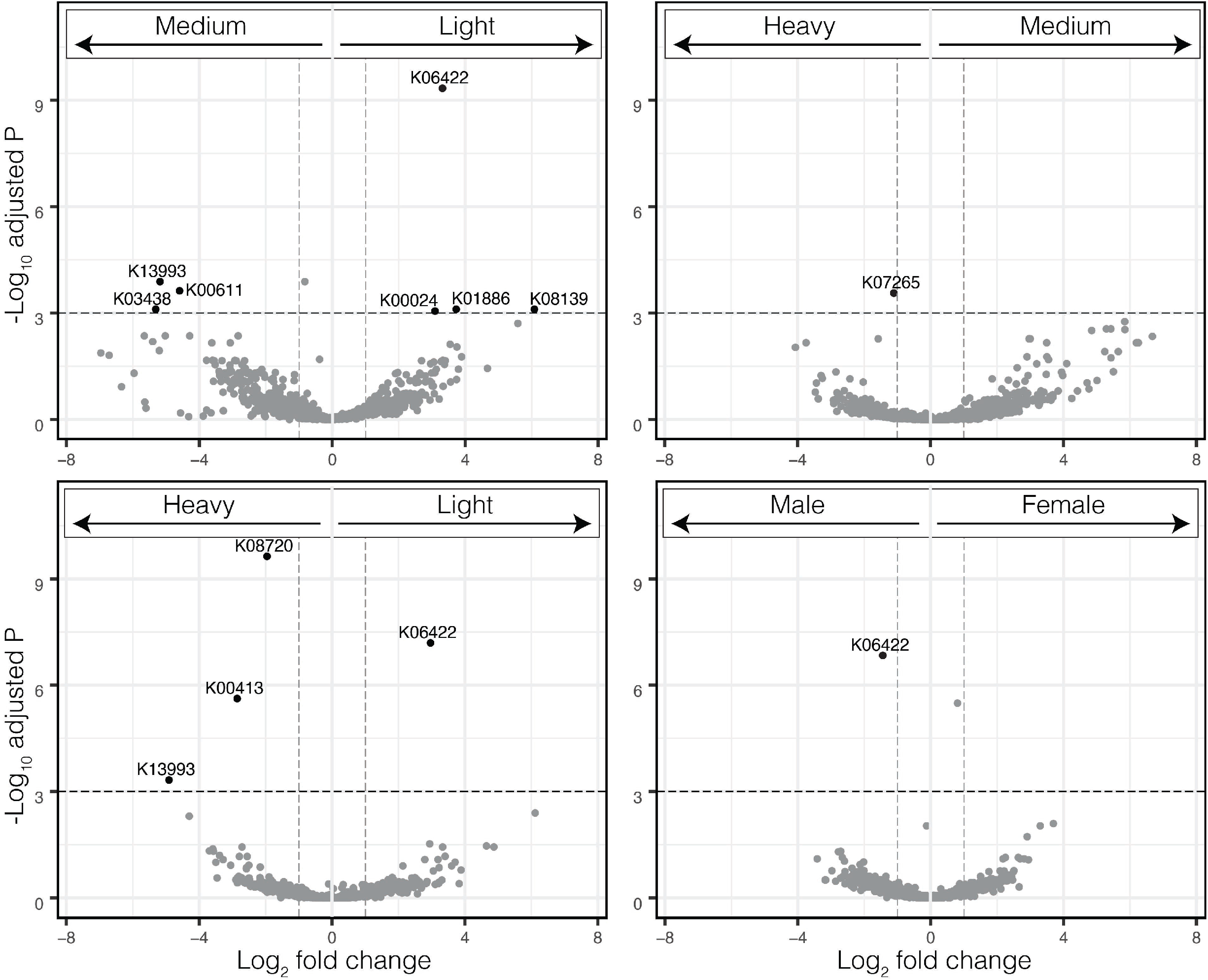
Volcano plots showing differentially expressed genes in metatranscriptomes from different weight classes and sexes of Ruddy Turnstones. The horizontal lines represent *p*=0.001. Genes that are differentially over or under expressed are identified with corresponding KEGG IDs.

Expression of three polyunsaturated fatty acids were significantly associated with weight (Fig. 6): linoleic acid metabolism (R^2^=0.19, *p=*0.024), alpha-linoleic acid metabolism (R^2^=0.23, *p=*0.014), arachidonic acid metabolism (R^2^=0.39, *p=*0.001). Biosynthesis of unsaturated fatty acids was not significantly associated with weight (R^2^=0.03, *p=*0.216).

**Figure 6.**
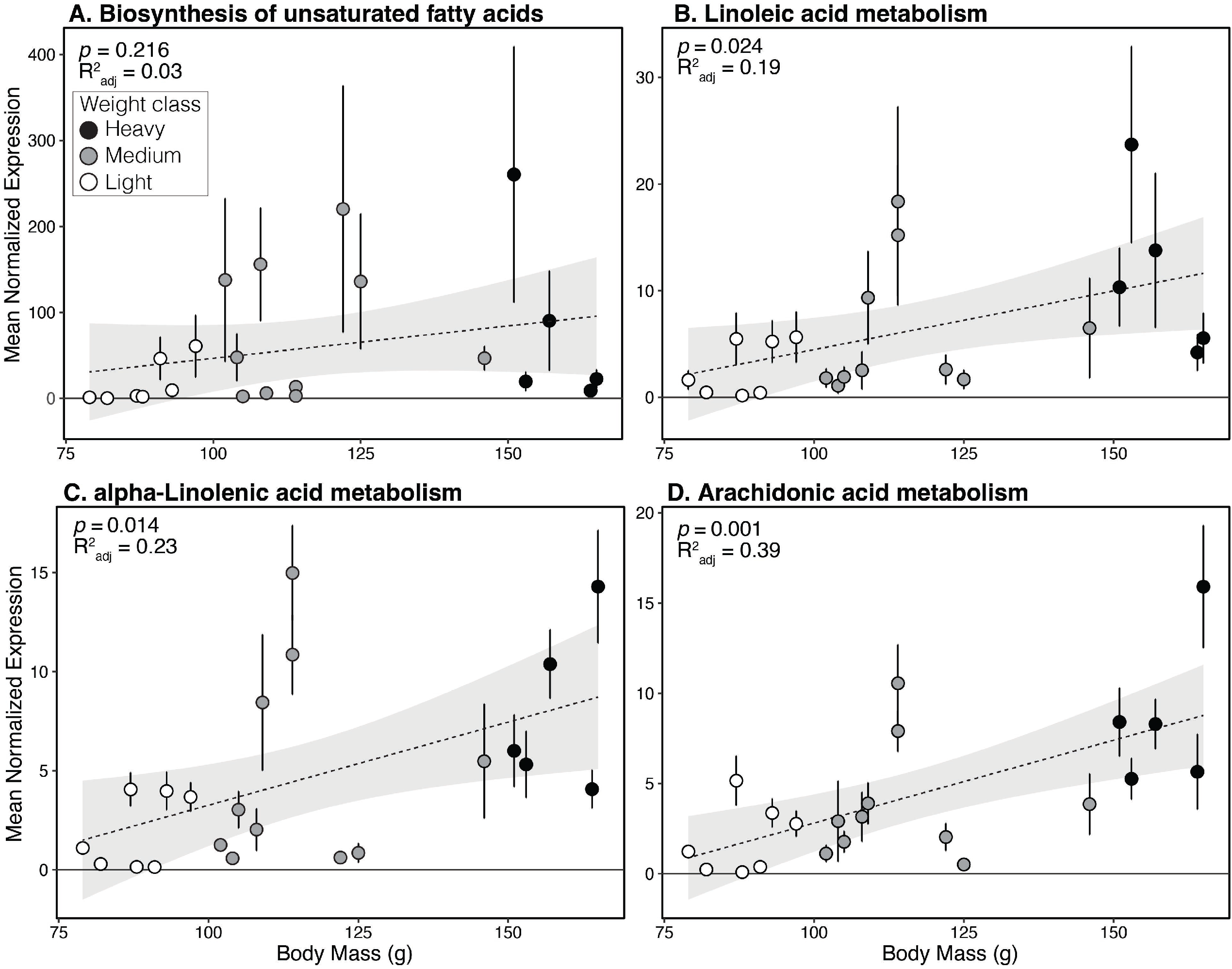
Linear regression (dotted line with gray 95% confidence intervals) of mean normalized expression of (A) Biosynthesis of unsaturated fatty acids, (B) Linoleic acid metabolism, (C) alpha-Linoleic acid metabolism, (D) Arachidonic acid metabolism. Error bars represent standard errors.

A clustering analysis was performed to group genes with shared expression patterns together, resulting in four clusters at *p<*0.05 (Fig. 7, Supplemental Table S4). Group 1 (N=130 genes) and Group 3 (N=76 genes) had light birds with contrasting expression patterns to the medium and heavy birds. Group 2 (N=9 genes) and Group 4 (N=22 genes) had medium weight birds as the highest or lowest expression group, respectively. All genes, regardless of significance, were analyzed as well and resulted in the same four expression patterns.

**Figure 7.**
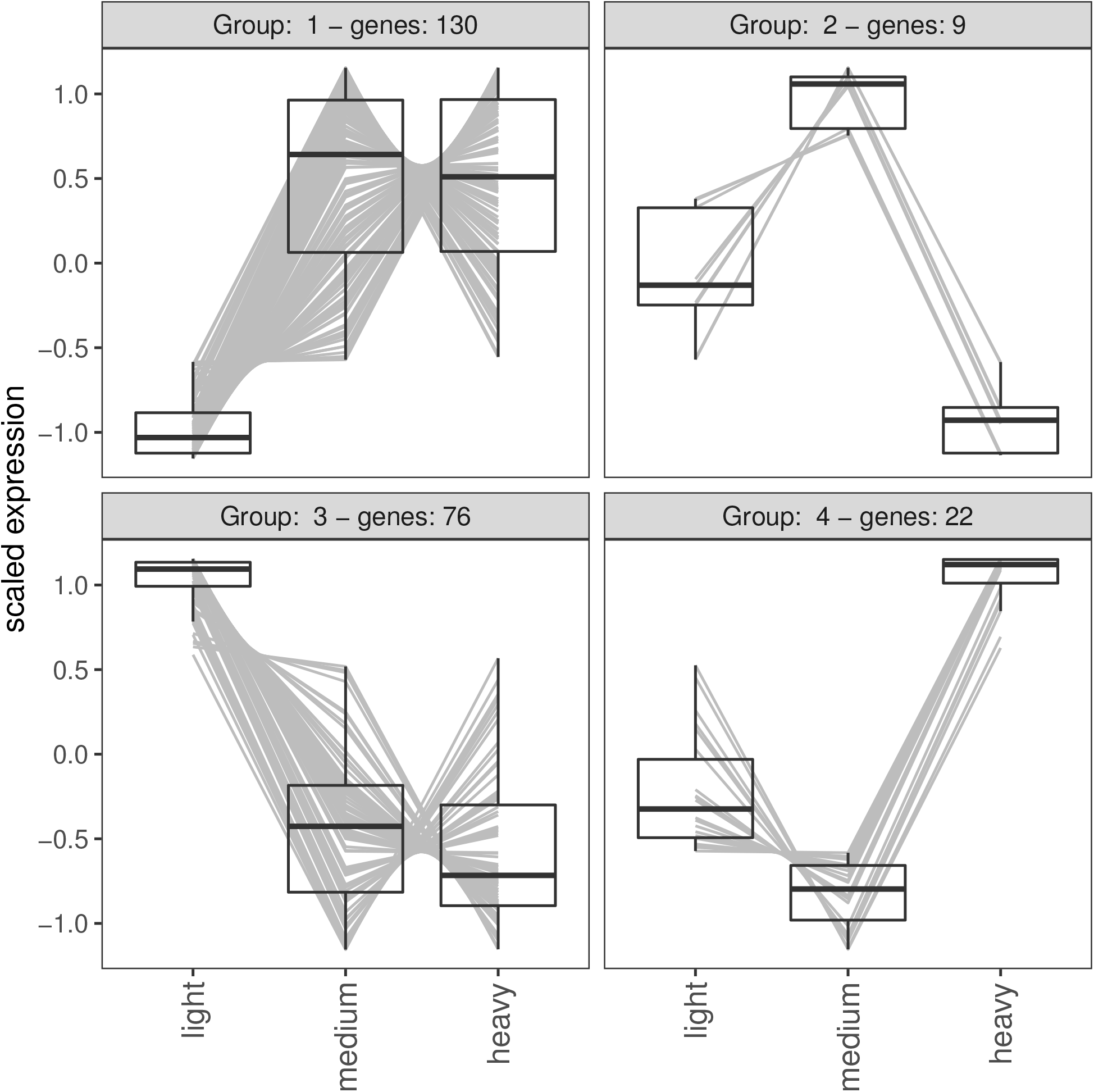
Clusters of genes with significantly similar expression across the weight classes (alpha=0.05). Y-axis represents Z-score. Positive values are upregulated compared to the average Z-score and vice versa for negative values.

## DISCUSSION

The microbiome is intertwined with host health, behavior and fitness and the microbiota can play a key role in weight gain in model organisms and humans [11, 58]. Shorebirds participate in extreme foraging behavior and rapid weight gain at stopovers during long distance migrations. Examining extreme behaviors or physiological processes may provide novel insight into the limits of how microbes facilitate vertebrate biology, and this has potential applied benefits to human health. In a broader sense, many animals experience periods of rapid or sustained weight gain (e.g., before migration or hibernation) or periods of weight loss, fasting or starvation that must be recovered from. Here, we have shown in a system with a uniform diet at geographically close sampling sites, that as the host’s body undergoes rapid weight gain, the taxonomic composition of the fecal microbiome changes in tandem with the host, as do some of the functional capabilities of the microbiome.

Migration causes a significant disturbance to a bird’s physiology and homeostasis; individual migrants can lose diversity of the microbiome during flight [59] and a corresponding successional recovery of the microbiota makes intuitive sense. Ruddy Turnstones gain an average of 50% body weight during stopover at Delaware Bay; therefore, the weight of a bird is a proxy for how long it has been at stopover. The three weight classes, light, medium and heavy, form significantly distinct clusters in our beta diversity ordinations based on 16S taxonomic composition and the medium weight birds are generally distributed between the low and heavy clusters (Fig. 2). This pattern implies successional change in the taxonomic composition of the fecal microbiome during stopover.

We used two methods to identify patterns within the taxa of the microbiomes: differential abundance analysis and ANOVA of the most abundant genera within the most abundant phyla. Succession within the microbiota may be further corroborated by the differentially abundant genera found across multiple pairwise tests (Fig. 3, Table S1). Of particular note is that the genera *Helicobacter, Catellicococcus* and *Campylobacter* are known bird gut-associated bacteria [60, 61] and these are more abundant in the light (or light and medium) birds. One hypothesis is that as the birds lose weight mid-migration, the bacteria that have co-evolved to live with the birds are in the mucus that remains in the gut while all transient or less well adapted bacteria are flushed from the system. As they fatten, environmental (e.g., *Maribacter, Denitromonas*), and fatty acid producing marine bacteria (e.g., *Vibrio, Shewanella, Photobacterium*, [62]) may flourish.

These taxonomic results contradicted our expectation that taxa from the Firmicutes would increase as birds gained weight, since Firmicutes have been associated with obesity or weight gain in mammals (but see [12]). The most abundant Firmicutes in the shorebirds were not significantly different between the weight classes (Fig. 3A). The genera *Flavobacterium* (Phylum: Flavobacteria) was significantly higher in heavy birds when compared to medium birds and *Vibrio* (Phylum: Proteobacteria) were significantly higher in the heavy birds when compared to both the light and medium birds (Fig. 3). Succession in microbiomes is seen in many vertebrate systems (e.g., [24, 63]) and occurs after periods of microbiome disturbance (e.g., antibiotic treatment). Our birds’ alpha diversity statistics trended upward the heavier they got, perhaps supporting a successional recovery of the microbiota. Continued investigation into the stability and resilience of microbiomes post-migration, and especially across years, would tell us how stable the shorebird microbiome is, on both short and long-term scales.

Sampling site was statistically associated with beta diversity in our 16S rRNA analyses, although with an effect size approximately one quarter of the weight class. We also detected a pattern between bird weight and the three sampling sites (Fig. S2). We hypothesize this is because sampling sites vary in food quality and better-quality sites are defended by larger birds. Broad scale associations between (food) quality of the sampling site and weight of the birds was observed, with lighter birds more frequently found on the lower quality sites, and heavier birds on the higher quality sites [64, 65].

As a community, and in contrast to the patterns displayed in taxonomic beta diversity (Fig. 2B), the medium weight functional communities were not obviously intermediate to the light and heavy weight categories (Fig. 4), although weight class was significantly associated with the variation of the samples (R^2^=0.22, *p=*0.02). To identify genes that may be associated with weight or weight gain, we used multiple methods: an unbiased pairwise significance test (Fig. 5), a cluster analysis (Fig. 7), and by specifically looking at pathways involved in polyunsaturated fatty acid biosynthesis (Fig. 6).

Using the pairwise significance test, most functions were not significantly different between the weight classes; only 10 functions (KEGG IDs, Table S3) rose above the significance threshold of *p<*0.001. Of these 10 functions, K06422 showed up most, as significantly higher in light birds (when compared to both medium and heavy birds) and as the only significant difference between males and females (higher in males). K06422 is associated with *sspE*, a small acid-soluble spore protein. As sporulation is a response to starvation in some bacteria, it may be that this protein is overrepresented in the light birds because they are underweight, and the microbiota had entered a stress/starvation response. The other functions associated with the light birds, K01886 and K08139 are also generally associated with cell growth and metabolism. The functions associated with the medium weight birds were all potentially associated with proteins (Arginine biosynthesis, ribosome biogenesis and heat shock proteins). K13993 is also associated with tissue remodeling and is significantly higher in both the medium and heavy birds when compared to light. This result seems counterintuitive as fattening shorebirds are known to switch from protein recovery post-migration to fat deposition around medium weight [5]. However, bacteria are known to digest dietary protein to produce secondary metabolites, such as amino acids, which could play a thus far unknown role in shorebird fattening [66]. Additionally, some tissues and organs are grown throughout the stopover period in other shorebirds [67] and may influence the microbiota.

The pathways identified in the heavy birds are more diverse than the previous groups. They include functions that refer to organismal systems and human diseases (K00413) and structural proteins (K07625). Of particular note is K08720, which is associated with *Vibrio* biofilm formation, and may specifically involve iron balance [68]. *Vibrio* was one of the taxa that was associated with heavier birds and is known to be a main microbial producer of polyunsaturated fatty acids [62]. *Vibrio* are also found in the horseshoe crab microbiome [69], so the increase could be due to dietary intake and benefit the host. Hosts could also be internally filtering for *Vibrio*, or both processes could happen in tandem.

The clustering analysis that grouped genes based on shared expression patterns detected four clusters. The 237 genes within these four clusters may contain interesting targets for groups of genes that perform together and differentially depending on bird weight. Since birds go through a period of protein recovery and immune suppression when they first land [70], genes in Groups 1 and 3 affected by those processes. The immediate arrival of birds also returns the microbes to a state of non-starvation, and genes in Group 3 could be many of those involved in cell growth and division. Conversely, and if the birds do benefit from the metabolites or products of the microbiota, as medium and heavy birds are prioritizing weight gain, the genes in Groups 1 and 3 may be downregulated and upregulated for fat deposition, respectively. The genes in Groups 2 and 4 could be those responding to the immediate recovery of the microbiota and the bird’s final preparations for flight; shorebirds both gain and lose particular muscles and tissues in response to and preparation for flight, and how the microbiota respond to those changes requires further investigation.

We investigated an *a priori* hypothesis that polyunsaturated fatty acid biosynthesis would increase as the birds fatten. “Essential” fatty acids are those an animal needs but cannot produce; the diet of horseshoe crab eggs provide the essential fatty acids birds need and PUFAs in particular are an extremely efficient way to store energy that shorebirds use to power their migrations. Alpha-Linoleic acid is an n-3 PUFA, whereas Lineoleic acid and Arachidonic acid are n-6 PUFAs. These different categories of PUFA can have multiple and antagonistic effects [71]. We hypothesize that the microbiome may also be producing essential fatty acids for the birds during weight gain. This unconventional hypothesis would require tracking experiments to confirm, but in our analyses, all three PUFAs (Linoleic acid, alpha-Linolenic acid and Arachidonic acid) significantly increased as the birds gained weight (Fig. 6). Is the increase of PUFAs simply a reflection of the microbiome rebounding and being a bigger collection of microbes? General biosynthesis of unsaturated fatty acids did not significantly increase with weight gain, indicating possible opposite patterns in other unsaturated FAs. A next step in our study is to investigate the full spectrum of fatty acids to identify patterns in abundance with weight change and pursue mechanistic explanations.

## DATA AVAILABILITY

Sequence and metadata will be made available on Figshare and the NCBI SRA upon acceptance.

## ACKNOWLEDGMENTS

We thank the Delaware Division of Fish and Wildlife and the Delaware Shorebird Project for allowing us to collect samples under their permits. This work was supported by the University of Connecticut [start-up funding to SMH]. Fieldwork of this project was funded, in part, through a grant from the United States Fish & Wildlife Service’s State Wildlife Grant Program. This work does not represent the opinions of the State of Delaware, Delaware Department of Natural Resources & Environmental Control or Delaware Division of Fish & Wildlife. Our work was conducted on the traditional and unceded land and territories of the Lenape Mohegan, Schaghticoke, Mashantucket Pequot, Eastern Pequot, Golden Hill Paugussett, and Nipmuc Peoples (CT), and the and Nanticoke Nations (DE).

## ETHICS STATEMENT

Samples were collected with permission from the Delaware Division of Fish and Wildlife-Department of Natural Resources and Environmental Control [2018-WSC-031 to KG], and the Federal Bird Banding Permit [23332 to Delaware Division of Fish and Wildlife-Department of Natural Resources and Environmental Control].

## SUPPLEMENTARY INFORMATION

**Supplemental Table S1**. All ANOVA p-values for detection of significant changes within genera across weight classes.

**Supplemental Table S2**. All genera identified as significantly overexpressed in the pairwise DESeq differential abundance analyses using amplicon data. Taxa that were detected in more than one pairwise comparison are bolded and on the same line of the table; the weight class (WC) in which the genus was overrepresented is shown and also denoted by color (yellow = Light, green = Medium, orange = Heavy).

**Supplemental Table S3**. All KEGG ID information for the functions identified as significant in the differential expression analysis.

**Supplemental Table S4**. All KEGG ID information for the functions identified as significant in the clustering analysis.

**Supplemental Figure S1**. Relationship between body weight and geographic sampling site.

**Supplemental Figure S2**. Individual level relative abundances of bacterial phyla, using 16S rRNA amplicon data.

